# Differential cross-reactivity to the influenza B virus haemagglutinin underpins lineage-specific susceptibility between birth cohorts

**DOI:** 10.1101/2023.08.25.554879

**Authors:** Peta Edler, Lara S.U. Schwab, Malet Aban, Michelle Wille, Natalie Spirason, Yi-Mo Deng, Michael A. Carlock, Ted M. Ross, Jennifer A. Juno, Steve Rockman, Adam K. Wheatley, Stephen J. Kent, Ian G. Barr, David J. Price, Marios Koutsakos

## Abstract

Influenza exposures early in life are believed to shape future susceptibility to influenza infections by imprinting immunological biases that engender differential cross-reactivity to future influenza viruses, but direct serological evidence linked to susceptibility is limited. We analysed hemagglutination-inhibition titres in 1451 cross-sectional samples collected between 1992-2020, from individuals born between 1917-2008, against influenza B virus (IBV) isolates from 1940-2021, including ‘future’ isolates that circulated after sample collection. We demonstrate that immunological biases are conferred by early life IBV infection and result in lineage-specific cross-reactivity of a birth cohort towards future IBV isolates. This translates into differential estimates of susceptibility between birth cohorts towards the two IBV antigenic lineages, explaining lineage-specific age distributions of observed medically attended IBV infections. Our data bridge a critical gap between early life exposure, cross-reactivity, and influenza epidemiology and identify a plausible model to further dissect the interplay between host immunity, viral evolution and epidemiology.

## Introduction

Influenza epidemiology is characterised by strong biases in susceptibility that cluster among individuals born within discrete year ranges (commonly referred to as birth cohorts)(1, 2). In this context, birth year can be used as a proxy of earliest influenza exposure, with individuals in the same birth cohort sharing a common earliest exposure to a specific influenza subtype, lineage or antigenic cluster (2). It has been proposed that these early life exposures establish immunological biases which modulate susceptibility to antigenically related influenza viruses encountered later in life, an effect termed ‘antigenic or immunological imprinting’ (3, 4). Such imprinting was initially proposed to explain differential protection against severe and fatal disease following infection with different avian influenza A viruses (IAV) (3, 4), but has since been applied to explain differential susceptibility of birth cohorts to medically attended disease by seasonal IAV subtypes or by influenza B virus (IBV) antigenic lineages (5, 6, 7, 8).

The association between earliest life exposure and protection later in life has been inferred from statistical models that reconstruct probabilistic exposure histories based on historical patterns of virus circulation, and retrospectively associate the most likely earliest exposure of a birth cohort to their observed susceptibility (3, 5, 6, 7). The mechanistic basis for the conferred protection has been proposed to lie within biases of HA-specific antibodies (2, 3). While these statistical models rarely incorporate immunological measurements, this notion is supported by analyses showing that individuals maintain the highest serological antibody titres against HAs from influenza virus isolates that circulated during their early childhood (9, 10). However, these studies analyse serological reactivity towards past and contemporaneous viruses, but not cross-reactivity to future unencountered and antigenically drifted variants. While it is assumed that antibody cross-reactivity to such future strains will be similarly impacted by imprinting and translate into differential susceptibility, there is little direct evidence to date to support these assumptions.

We reasoned that IBV constitutes a tractable model to test these assumptions due to the discrete circulation patterns of IBV antigenic lineages between 1940-2000 (11, 12), and the distinct patterns of birth cohort- and lineage-specific epidemiology observed between 2000-2020 (7, 13, 14, 15, 16, 17). In the first 3 decades since its discovery in 1940, IBV circulated as a monophyletic lineage, which we refer to as “Ancestral”. In the early 1970s, a lineage (later designated as B/Victoria) appeared that was antigenically distinct from Ancestral viruses (11, 12). In the 1980s, a second antigenically distinct lineage emerged (designated as B/Yamagata)(12) which predominated in the 1990s, with B/Victoria remaining confined to low levels within Asia (11). Subsequently, the B/Victoria lineage re-emerged globally, and the two lineages co-circulated from the early 2000s until the putative extinction of B/Yamagata in 2020 (18, 19). During co-circulation of both B/Yamagata and B/Victoria viruses from 2000-2019, differential susceptibility of adults to medically attended disease was observed based on their birth cohort (7). During that time, individuals born in the 1980s and 1990s (adults in their 20s-40s) were relatively protected from medically attended infections caused by B/Yamagata, while the pre-1980 birth cohorts (adults in their 40s-70s) experienced limited medically attended infections by B/Victoria (7, 14, 15, 16, 17). It has been proposed that these epidemiological patterns of susceptibility may have resulted from immunological biases established their first life exposure to influenza B viruses (B/Victoria viruses before 1980 and B/Yamagata in the 1990s) (7).

To test this hypothesis and the assumptions underlying the imprinting model of influenza susceptibility, we analysed hemagglutination inhibition (HI) titres against IBV in individuals born over 9 decades. We show that HI titre differences towards future unencountered isolates translate into birth cohort-specific estimates of susceptibility and likely underpin the epidemiology observed for the two IBV lineages. Our findings thus provide strong immunological evidence that support the link of early life exposure as a major determinant in the differential protection against influenza B disease.

## Results

### Dissecting HI reactivity profiles against IBV antigenic lineages across birth cohorts

To assess for biases in serological reactivity against IBV across birth cohorts, we analysed baseline (prior to any annual vaccination) HI measurements from three separate datasets. Firstly, we identified 322 individuals born between 1917-2008 who were cross-sectionally sampled between 1992-2020 at an age of 1-83 years old (Figure S1A). Serum or plasma samples were titrated in HI assays against 19 IBV isolates collected from 1940-2021 (Figure 1A). We refer to this dataset of 6118 HI titrations as the main dataset of the study. Isolates circulating prior to 1980 were chosen based on availability, while viruses from 1980 onwards were WHO nominated vaccine strains. Recently, B/Hong Kong/05/1972 was identified as the earliest isolate belonging to the B/Victoria lineage, demonstrating the emergence of B/Victoria in the 1970s (12). In support of this, we observed strong correlations (r = 0.6-0.78) (Figure S2A) between HI titres to B/Hong Kong/05/1972 and all 8 B/Victoria isolates studied, but none of the 5 B/Yamagata isolates studied. The similarity of B/Hong Kong/05/1972 with B/Victoria isolates is consistent with the presence of the B/Victoria-defining 150N mutation (12), although we note that B/Hong Kong/05/1972 titres were also correlated (r = 0.61-0.73) with titres to Ancestral B isolates, possibly reflecting the conservation of other epitopes between B/Hong Kong/05/1972 and Ancestral isolates. Similarly, we found strong correlations (r = 0.586-0.68) between HI titres to B/Austria/1359417/2021 (B/Victoria) and Ancestral B isolates, supporting antigenic similarity between recent B/Victoria isolates and ancestral viruses, consistent with previous reports and the presence of the lineage defining 150K mutation(12). We further observed these antigenic similarity patterns using five previously published monoclonal antibodies (mAbs) (20, 21, 22) (Figure S2B). As we included serum samples obtained as early as 1992 and viruses isolated as late as 2021, 27.5% of the HI measurements were against isolates that circulated after sample collection, which we refer to as ‘unencountered future viruses’ (Figure S1B). These future isolates represent antigenic variants circulating between 2000-2022 during which differential susceptibility to the two IBV lineages has been observed.

**Figure 1.**
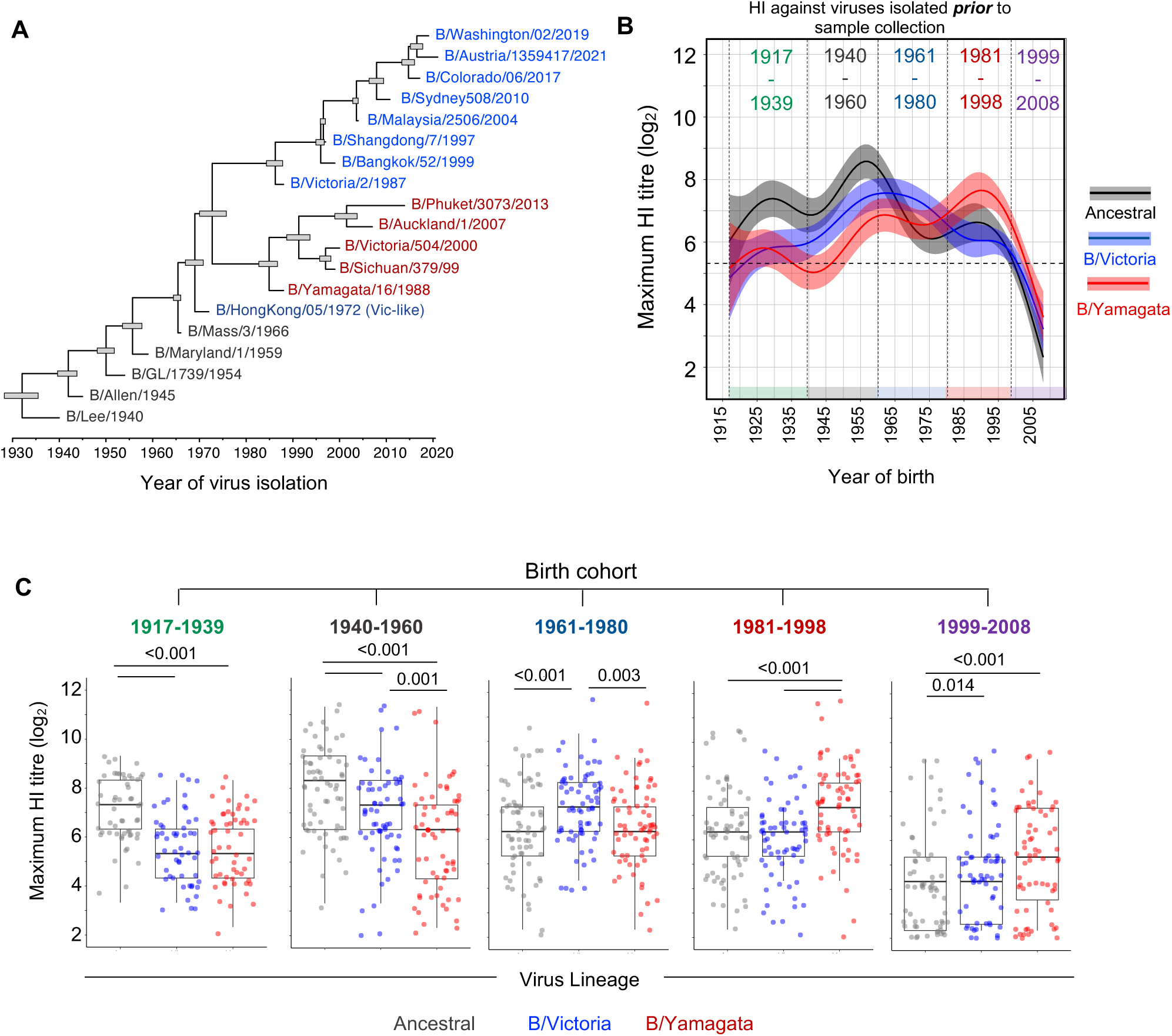
Antibody reactivity against past isolates from different lineages varies by birth year. **(A)** Time structured phylogenetic tree of the HA nucleotide sequences for the 19 selected viruses used for serological analysis for the main dataset. Node bars correspond to the 95% highest posterior density of node height. Scale bar represents time, in years. **(B)** Maximum HI titres against the three IBV lineages spanning 1940 and 2020, across birth years. The lines represent estimates using generalized additive models (GAMs) with 95% CI (n=322 individuals from the main datasets). Only titrations of viruses isolated prior to sample collection are included. **(C)** Box-plots of paired maximum HI titres to each lineage for individuals within each birth cohort (n=54 for 1917-1939, n=65 for 1940-1960, n=70 for 1961-1980, n=67 for 1981-1998 and n=66 for 1999-2008). P-values were generated from a Friedman’s test with Dunn’s correction for multiple comparisons.

The main dataset was complemented by an independent, previously reported (23) HI titre dataset of 85 individuals aged 24-80 years old, sampled in 2014 and titrated against 16 IBV isolates from 1940-2017 (Figure S1C), which we refer to as the ‘Carlock et al’ dataset. Finally, we compiled cross-sectional HI data from 1044 individuals from annual serology testing at the WHO Collaborating Centre for Reference and Research on Influenza (WHOCCRRI) in Melbourne between 2000 and 2020 that were assayed against one or two contemporary isolates (overall 5 B/Victoria isolates and 7 B/Yamagata isolates) (Figure S1D), which we refer to as the ‘WHOCCRRI’ dataset. The two supplementary datasets were used to validate observations from the main dataset where appropriate.

Despite analysing samples collected over a 29-year period, we did not observe a bias in HI reactivity towards viruses circulating closest to the year of sample collection (compiled data from all 3 datasets, Figure S2C). Additionally, the earliest sampled cohort (1992) did not have lower HI titres to the more distant future unencountered isolates compared to more recently sampled cohorts (Figure S2D). Due to availability of viral isolates, all datasets included HI measurements against some egg-propagated isolates. These comprised 47.4% of isolates in the main dataset and were limited to viral isolates prior to 1999. HI titres against a recent B/Victoria or B/Yamagata isolates were approximately 2-fold higher against egg isolates than paired HI titres against their equivalent cell isolates (Figure S2E) but the two measurements were strongly correlated (Figure S2F). HI titres were also strongly correlated with microneutralization titres against live IBV (Figure S2G), verifying the utility of HI as a surrogate for neutralising antibodies. We therefore analysed the main dataset and two additional datasets to determine if there were immunological biases that may explain the observed differences in IBV epidemiology.

### HI reactivity against past isolates from different lineages varies by birth year

We firstly considered the effects of birth year on HI titres against each past isolate (circulating prior to sample collection) from each IBV lineage (ancestral, B/Victoria and B/Yamagata) (Figure S3A). We also estimated the mean HI titre against year of birth for each lineage by fitting a generalised additive model (GAM) to pooled HI titres for all past isolates, with participant ID as a random effect, to account for the individual-level variability (Figure S3B). HI titres against the Ancestral isolates were highest in individuals born around the 1950s and 1960s than in other birth years. Similarly, HI titres to B/Victoria were highest in individuals born around the 1960s and 1970s than in other birth years, while HI titres against B/Yamagata were highest in individuals born in the 1980s and 1990s than in other birth years. These patterns were consistent across the main and two supplementary datasets (Figure S3). We note that HI titres to B/Lee/1940 were uniformly high in all adults across both datasets. To further assess how birth year impacts reactivity towards the three lineages within each individual, we determined the maximum detectable HI titre for each lineage per individual from the main dataset (Figure 1B). This recapitulated the effects of birth year on HI towards each lineage but further indicated a hierarchy between lineages that varies by birth year.

To facilitate analysis of HI by birth year, lineage and other variables, we grouped individuals into five birth cohorts based on previous analysis of probabilistic infection histories to IBV (7) as well as the antigenic phenotypes of viruses circulating between 1959-1990 (12) (see Methods section for details). Specifically, we considered the 1917-1939 and the 1940-1960 birth cohorts with earliest life exposure to IBV from the Ancestral lineage (the distinction being the lack of IBV isolates prior to the discovery of IBV in 1940); the 1961-1980 birth cohort, with most-likely earliest life exposure to early B/Victoria viruses of that time (12); the 1981-1998 birth cohort, with most likely earliest life exposure to B/Yamagata; and the 1999-2008 birth cohort with mixed early life exposures to either B/Yamagata or B/Victoria. Although the selected birth year boundaries are somewhat arbitrary, the five birth cohorts recapitulate the variability of maximum detectable HI titres by birth year (Figure 1B, C). Indeed, by comparing paired maximum HI titres to past isolates for each lineage within each individual (Figure 1C, descriptive statistics provided in Table S1), we found that on average the 1917-1939 had higher HI titres to Ancestral isolates than to other lineages; the 1940-1960 birth cohorts had higher HI titres to Ancestral isolates than to other lineages and higher HI to B/Victoria than B/Yamagata; the 1961-1980 birth cohort had higher HI titres to B/Victoria isolates than to other lineages; the 1980-1998 birth cohort had higher HI titres to B/Yamagata isolates than to the other lineages. The 1999-2008 birth cohort also had higher HI titres to B/Yamagata, but it had the lowest HI titres overall compared to other birth cohorts likely due to their younger age. We observed overall similar patterns when comparing HI titres between birth cohorts for each isolate separately, consistently across the main and two supplementary datasets (Figure S4A-C, descriptive statistics provided in Table S2-4). We did however also observe some differences between specific isolates. For instance, HI titres to B/Yamagata/16/1988 were higher in the 1961-1980 cohort than the 1981-1998 cohort. Additionally, titres to more recent B/Victoria isolates were not different between the 1961-1980 and the 1981-1998 cohorts (e.g. B/Malaysia/2506/2004), or across all birth cohorts (e.g. B/Washington/02/2019). Furthermore, the 1917-1939 birth cohort had the highest HI titres to B/Allen/1945 compared to other birth cohorts, while the 1940-1960 cohort had highest HI to B/GL/1954, B/Maryland/1959 and B/Mass/1966 (Figure S4A). These patterns suggest that there might be biases towards IBV lineages circulating in early life, as well as towards different isolates within each lineage. Overall, we conclude that individuals from different birth cohorts demonstrate a specific bias in HI reactivity towards one of the three IBV antigenic lineages, compared to other lineages and other birth cohorts.

### Early life exposure and antigenic seniority shape birth cohort-specific HI reactivity against IBV lineages

We next assessed how these biases were associated with early life exposure. Firstly, considering all past isolates, we determined the age of individuals at the time of circulation of the isolate for which peak titres were observed (Figure S5A). Across lineages and birth cohorts, average peak HI titres of each individual were observed for the isolate that circulated around birth. This could be further illustrated by analysing HI reactivity towards viruses isolated in different years for each birth cohort (Figure 2A, descriptive statistics provided in Table S5). Indeed, for each birth cohort we found an average peak in HI for isolates circulating during the years of birth of that cohort, or the subsequent 5-10 years. For instance, the 1961-1980 birth cohort on average highest titres towards B/Hong Kong/5/1972. We also noted that the 1940-1960 and 1961-1980 cohorts had nearly as high titres for B/Phuket/3073/2013 from the B/Yamagata lineage, which may reflect recent exposure of these cohorts to this lineage, although only 20% of individuals in these cohorts were sampled after B/Phuket-like viruses circulated. Nonetheless, these data demonstrate that individuals can maintain relatively high HI titres to isolates encountered early in life, even if that was multiple decades prior to sample collection.

**Figure 2.**
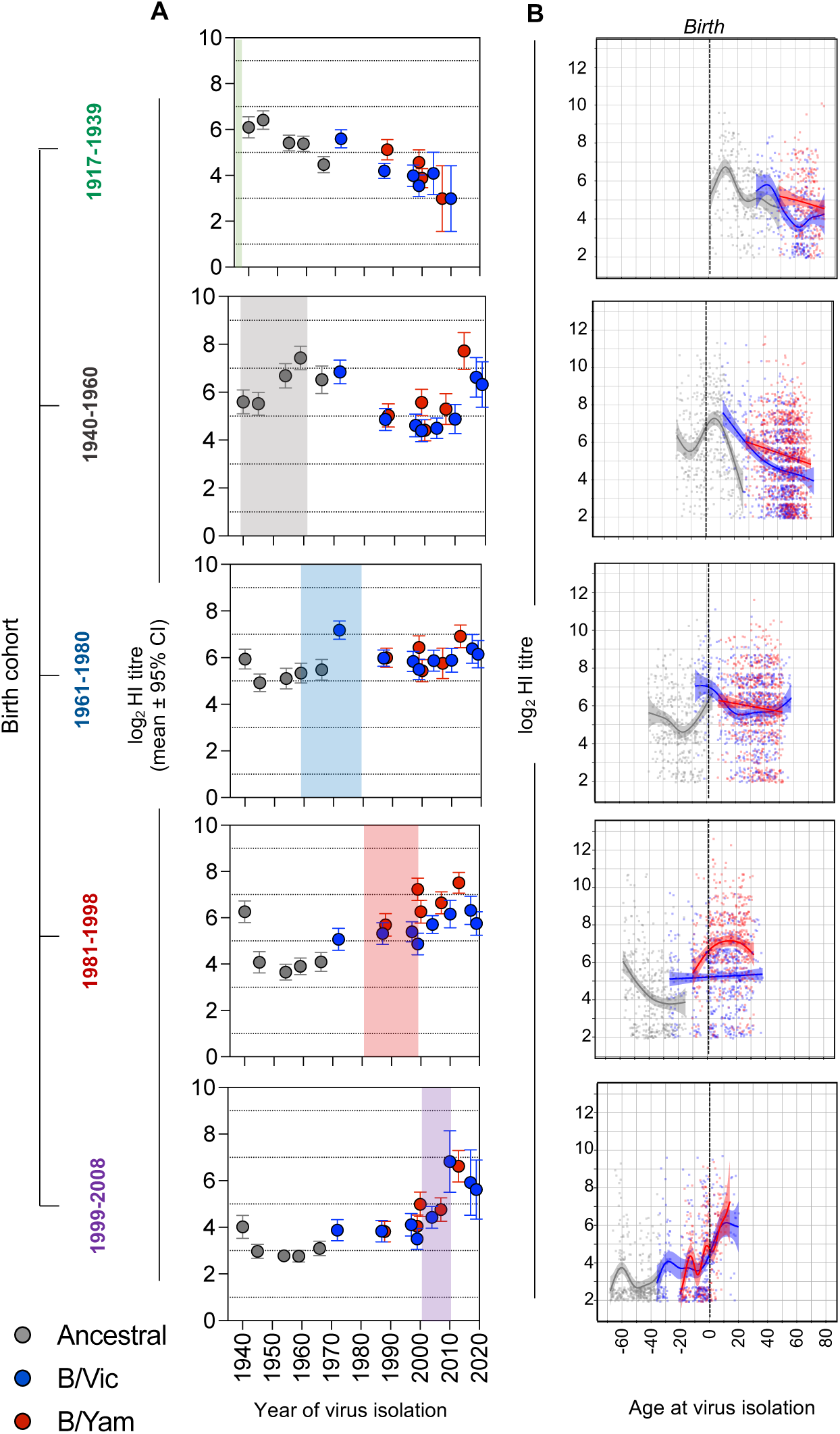
Early life exposures and antigenic seniority explain birth cohort-specific reactivity to different lineages. **(A)** Mean log_2_ HI titres to individual isolates for each birth cohort against year of virus isolation. Virus isolates are color coded by IBV lineage. The shade area represents the years of birth of that cohort. **(B)** Antibody titres relative to the age of the participant at the time of virus isolation for each lineage separated by birth cohort. The lines represent estimates using generalized additive models (GAMs) with 95% CI, accounting for repeated measurements on each individual by specifying a random effect. Only titrations of viruses isolated prior to sample collection are included.

To further demonstrate that HI titres to IBV isolates encountered early in life are maintained at higher levels than titres to isolates encountered later in life (i.e., antigenic seniority)(10), we considered the age of individuals at the time of virus isolation for all available past isolates. HI titres were on average highest for past viruses that circulated in the first decade of life and decreased for isolates circulating later in life (Figure S5B). Maximum HI titres were observed for isolates circulating at ∼8 years of age, consistent with observations in antibody titres against H3N2 IAV (10). This was also observed in the supplementary dataset by Carlock et al. This 8-year window explains the patterns observed in Figure 2A where individuals had average peak HI for isolates circulating during the year of birth of that cohort, or the subsequent 5-10 years. When considering the age at sampling, HI titres increased for the first 2 decades of life, but remained relatively stable thereafter, until approximately the age of 60 when they started declining (Figure S5C). When we analysed HI titres against age at virus isolation separately for each birth cohort and lineage (Figure 2B), we observed that each birth cohort had highest titres to isolates from the lineage circulating around birth, supporting antigenic seniority at the level of antigenic lineages.

Collectively our analyses support the hypothesis that because infections in childhood have a long-lasting effect on the specificity of HI reactivity, the distinct patterns of IBV lineage circulation over the last 80 years have resulted in birth cohort-specific biases in HI reactivity against the three IBV lineages.

### HI cross-reactivity to future isolates from different IBV lineages varies by birth year

We next considered if the same HI reactivity biases extend to unencountered future isolates, which will likely be antigenically drifted. Since our dataset included titrations to IBV isolates that circulated after sample collection, we were able to assess HI titres to future unencountered isolates from both the B/Victoria lineage and the B/Yamagata lineage (Figure 3A and S6A-B). Specifically, we found that individuals born between 1981-1998 had on average higher HI titres against unencountered B/Yamagata isolates than other birth cohorts and 2-fold higher B/Yamagata-specific HI titres than towards future unencountered B/Victoria isolates. Conversely, imprinting of B/Victoria HI cross-reactivity was evident in earlier birth cohorts, specifically those born between 1940-1980. A similar pattern was observed for HI titres against B/Colorado/06/2017 in the Carlock et al dataset (Figure S6A-B).

**Figure 3.**
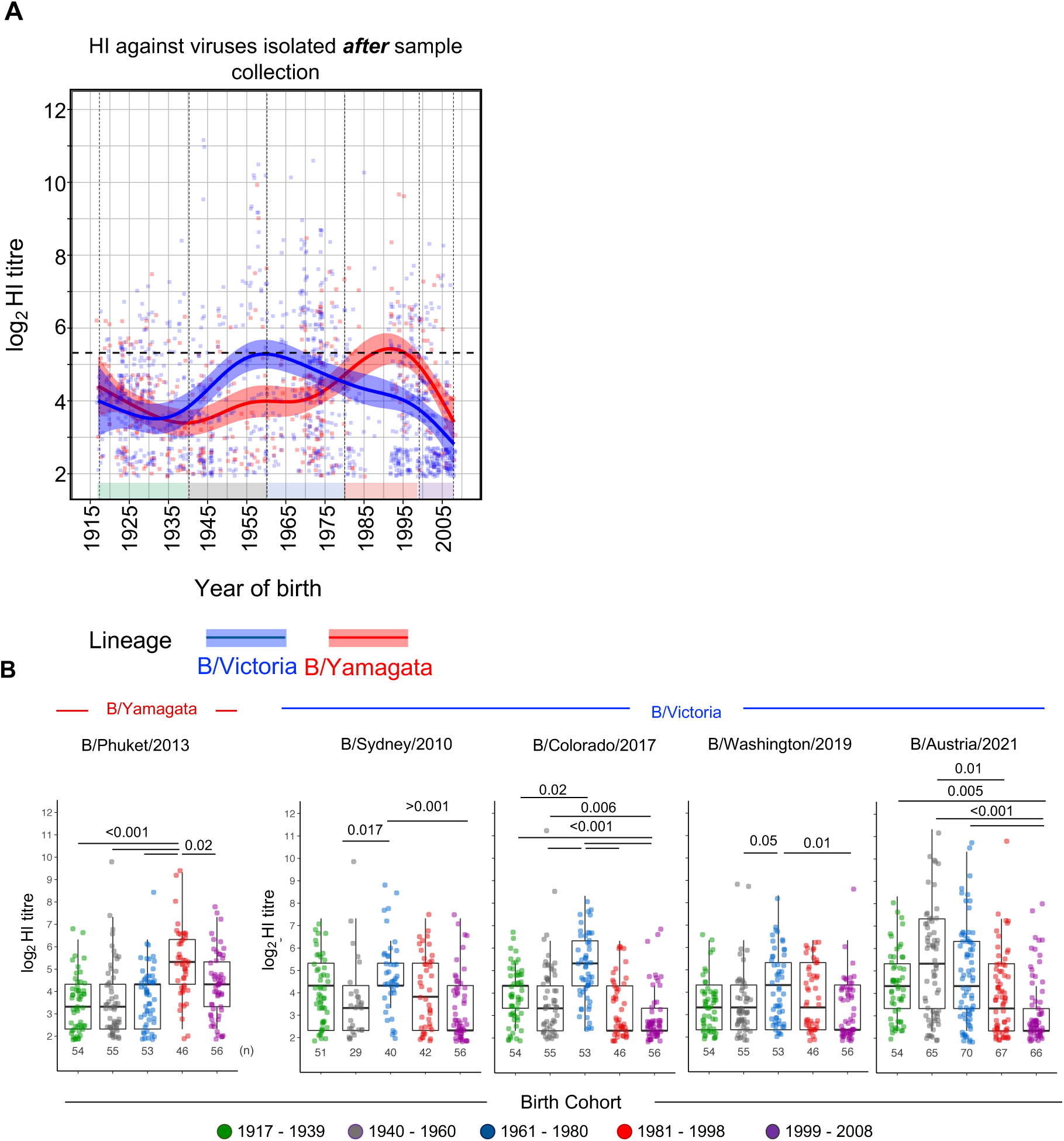
HI cross-reactivity towards unencountered future isolates from different IBV lineages varies by birth year. **(A)** Estimated mean HI titres against future unencountered isolates that circulated after sample collection, against the year of birth for each participant. The lines represent estimates using generalized additive models (GAMs) with 95% CI, accounting for repeated measurements on each individual by specifying a random effect. Dots show individual participants colored by birth cohort. **(B)** Box-plots of HI titres to specific future unencountered IBV isolates for each birth cohort. Samples size (n) is indicated at the bottom of each graph. P-values were generated from a Kruskal-Wallis test with Dunn’s correction for multiple comparisons.

Next, we performed a sensitivity analysis to determine whether the estimated mean HI titres for each lineage were consistent if any single isolate was excluded. Overall, estimated HI titres against future B/Yamagata or future B/Victoria were similar to the main analysis (Figure S6C). The only exception was exclusion of the B/Austria/1359417/2021 HI titres, which provided a B/Victoria HI estimate that was highest around 1970-1980 birth years, compared to a peak around 1960-1970 as seen in the primary analysis. Additional sensitivity analysis showed that the estimated HI titres were similar if cohorts of different sampling years from the main dataset were excluded (Figure S6D). The only exceptions were the effects of the 2020 cohort, whose only unencountered isolate was B/Austria/1359417/2021 thus recapitulating the effects of removing this isolate (Figure S6D); and the effects of the 2009 cohort, which exclusively comprised of samples from children (<18 years old), whose titres are overall lower than those of adults (Figure S5C). The patterns of birth cohort-specific HI cross-reactivity to future isolates were further evident when we analysed HI titres to individual isolates for which data were available across all five birth cohorts (Figure 3B, descriptive statistics provided in Table S6). Specifically, while individuals in the 1981-1998 cohort (B/Yamagata-imprinted) had on average higher HI titres to B/Phuket/3703/2013 (B/Yamagata), those in the 1961-1980 birth cohort (B/Victoria-imprinted) had higher HI titre against B/Sydney/805/2010 (B/Victoria), B/Colorado/06/2017 (B/Victoria) and B/Washington/02/2019 (B/Victoria) than other birth cohorts. In contrast, to these earlier B/Victoria viruses, the average highest HI titre against the B/Austria/1359417/2021 isolate were observed for the 1940-1960 birth cohort.

The HI results with of the B/Austria/1359417/2021 isolate are noteworthy as ferret anti-sera specific for Ancestral isolates show HI cross-reactivity to B/Austria/1359417/2021 but not earlier B/Victoria isolates likely due to the presence of the lineage-defining 150K mutation in Ancestral isolates and the B/Austria/1359417/2021 (12). This explains the HI reactivity towards the future unencountered B/Austria in the 1940-1960 birth cohort. These data suggest potentially distinct imprinting by Ancestral IBV viruses from 1940s-1960s and by early B/Victoria viruses from the 1970s. However, additional immunological and epidemiological analyses are required to investigate this observation and its implications. Nonetheless, our data overall support the idea that differential early life exposure to different IBV lineages between birth cohorts results in differential serological HI cross-reactivity towards future unencountered and antigenically drifted IBV isolates.

### Probability of infection with unencountered future isolates from different IBV lineages tracks with IBV epidemiology

To understand the effect of birth cohort-specific HI cross-reactivity on IBV susceptibility, we considered the previously established relationship between HI titres and protection from influenza virus infection (24). We simulated HI titres from the established GAMs, to maintain the correlation between HI titres to different viruses. Using the sero-protection curve by Coudeville et al. (24), we estimated the probability of infection for individuals born in different years corresponding to these HI titres.

When we estimated the probability of infection for previously circulating and possibly encountered isolates from each lineage (Figure 4A), we found that individuals born between 1960-1980 and between 1981-1998 had similar estimates for both B/Yamagata and B/Victoria. In contrast, the probability of infection for future unencountered isolates was birth cohort and lineage-specific (Figure 4B). Specifically, individuals born between ∼1981-1998 had the lowest estimated probability of infection with future B/Yamagata compared to other birth cohorts and to future B/Victoria. Conversely, individuals born between ∼1950-1980 had the lowest estimated probability of infection with future B/Victoria viruses compared to other birth cohorts and to future B/Yamagata viruses. Sensitivity analyses of the estimates of susceptibility (Figure S6E-F) were consistent with those of HI titres (Figure S6C-D). The estimated differences in susceptibility towards the two IBV lineages are highly consistent with the observed birth year distributions of B/Yamagata and B/Victoria medically attended cases observed globally between 2000-2022 (based on the number of sequences deposited on GISAID) (Figure 4C). We note, however, that for individuals born prior to 1950 we estimated a high probability of infection with B/Victoria, in contrast to the paucity of B/Victoria cases in that birth cohort. This may reflect that this birth cohort comprises a smaller fraction of the population, higher vaccination rates of this high-risk group, and/or the involvement of protective immune effectors other than HI antibodies in that group. Nevertheless, overall, this analysis strongly links HI cross-reactivity to future isolates with susceptibility and provides a potential immunological basis for the contrasting epidemiology of the two IBV lineages.

**Figure 4.**
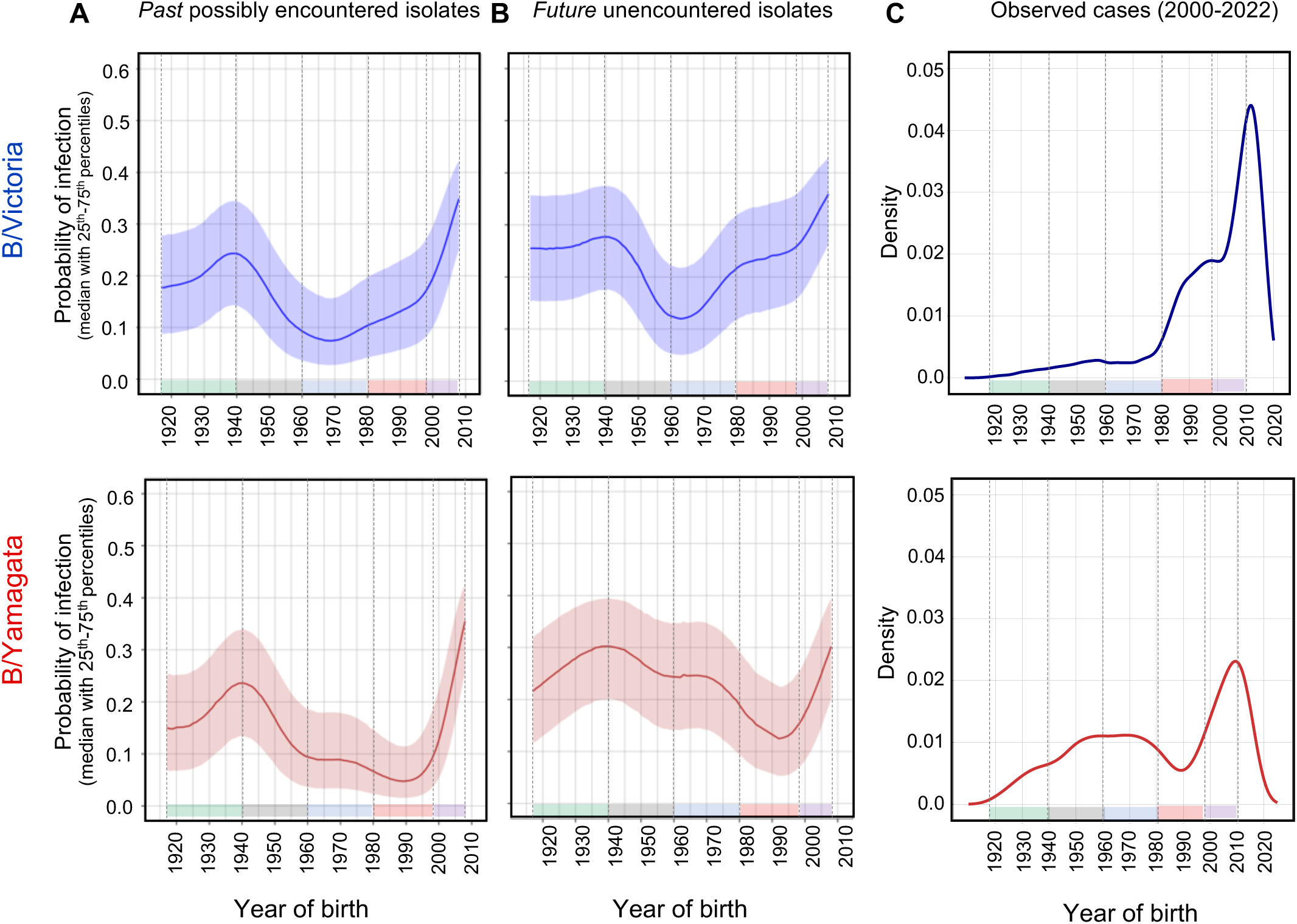
Probability of infection for future isolates from different IBV lineages varies by birth year. **(A)** Estimates of the probability of infection for past, possibly encountered, IBV isolates from each lineage by birth year. The probability of infection was estimated using the sero-protection curve by Coudeville et al.(24) and simulated HI titres from the established GAMs for past isolates for individuals born in different years. **(B)** Estimates of the probability of infection for future unencountered IBV isolates from each lineage by birth year. The probability of infection was estimated using the sero-protection curve by Coudeville et al.(24) and simulated HI titres from the established GAMs for future possible isolates for individuals born in different years. The median probability estimate is shown and the shaded areas represent the 25^th^ and 75^th^ percentiles. **(C)** Distribution of IBV cases by birth year observed between 2000-2022 group by lineage. Cases are based on the number of sequences deposited on GISAID.

## Discussion

The effects of influenza virus exposure early in life are considered a critical determinant of influenza susceptibility by establishing life-long biases in immunological reactivity to influenza viruses (1, 2). Such immunological biases have been well-established in cohorts with recent re-exposure to influenza A viruses (9, 25, 26, 27). However, a link between specific biases in baseline antibody titres and subsequent susceptibility has not been established for either IAV or IBV. Our analysis which includes cross-reactivity to future viruses, thus, provides an immunological basis linking early life exposure and susceptibility to influenza disease. This has substantial implications, not only for mitigating the clinical burden of IBV, but also by identifying IBV as a potential model to further dissect the interplay between host immunity, epidemiology and viral evolutionary dynamics.

The differential age distribution of medically attended infections by B/Yamagata and B/Victoria have been well established (7, 14, 15, 16, 17). This could either be explained by the combined lineage-specific usage of sialic acid receptors and their differential expression across age groups (15), or by birth-cohort effects of differential early life exposure (7, 14). Although based on our data we cannot exclude the contribution of sialic acid expression patterns, our immunological analyses strongly support a role for antibody imprinting by early life exposure to IBV as the driver of differential HI reactivity and subsequently the divergent epidemiology of the two IBV lineages. It will be important to determine these immunological biases can shape vaccine effectiveness, as has been described for the H1N1 and H3N2 vaccine components (28, 29, 30). What also remains unclear is whether subclinical/asymptomatic infections are similarly impacted by imprinting or if this only relates to medically attended infections (31). If IBV epidemiology has an immunological basis, immunological interventions early in life, including immunisation with an appropriate formulation of a universal IBV vaccine, may have a considerable impact in mitigating the clinical burden of IBV in the face of ongoing evolution.

Our study raises the question of how such biases in HI reactivity are maintained in a lineage-wide manner despite antigenic drift within each lineage (32, 33) and IBV re-infections (34). This is likely underpinned by the propensity of the immune system to recall cross-reactive memory B cells established by prior exposure, as has been described for other viruses (35, 36, 37). We hypothesize that re-exposure to the imprinted IBV lineage boosts antibodies to conserved epitopes including neutralising ones, while secondary exposure to a non-imprinted IBV lineage boosts antibodies to conserved but non-neutralising epitopes. Additionally, exposure to a non-imprinted IBV lineage will elicit neutralising antibodies to this secondary lineage but at levels lower than those of the imprinted lineage. While additional studies in IBV-infected or vaccinated cohorts are required to fully test this model, previously, IBV HA-specific monoclonal antibodies isolated from adults receiving trivalent influenza vaccine primarily showed either lineage-specific HI activity or cross-lineage binding without HI activity (20). Although some cross-lineage HI boosting in serum has been reported after vaccination (23, 38) or infection (39), this has been observed at acute timepoints after re-exposure, and the extent to which this is maintained long-term is unclear. Indeed, long-term antibody landscape analyses of HI to H3N2 viruses following vaccination or re-infection indicate that the broad recall of antibodies observed at the acute phase of re-exposure subsides with time, and the long-term immunological gain is focused on the re-exposure variant (9, 40, 41). It is important to determine if re-exposure to IBV similarly shapes HI reactivity landscapes across birth cohorts. This is particularly pertinent in considering how immunity towards the potentially extinct B/Yamagata lineage may wane or evolve over time, as our analysis indicates that certain birth cohorts could be susceptible to disease by B/Yamagata viruses should this lineage re-emerge. The lineage-wide HI reactivity observed for IBV suggests the existence of conserved antigenic sites enabling neutralisation within a given lineage, but most likely not shared between lineages. In support of this, human mAbs with pan-lineage HI activity have been previously isolated (20, 21) (Figure S2B). Characterising such conserved epitopes within different lineages, and strategies to boost responses towards them by immunisation could facilitate improvements in the efficacy of influenza vaccines against IBV in the face of ongoing antigenic drift. This, however, will require a detailed understanding of the IBV HA antigenic space and its molecular basis, which is currently limited (12, 15, 32, 33).

Our study has several limitations. As our analyses are based on cross-sectional samples, we cannot directly associate the HI titres within an individual to their actual protection from influenza. We have no knowledge of actual exposure histories (influenza infections or prior vaccinations) for individuals within our cohorts, which will vary. Nevertheless, the HI biases observed here are highly consistent with the infection histories inferred by statistical models (7). Thus, despite likely heterogeneity, immunological imprinting is still evident at the population level in different birth cohorts. In addition, we focused on HI titres and have not considered if other immunological factors that could contribute to protection, like virus microneutralization titres, HA stem-specific or NA-specific antibodies (42) similarly could vary by birth year. Furthermore, our estimates of susceptibility based on HI titres do not consider changes in serological profiles over the sampling period (1992-2020). However, sensitivity analyses did not identify biases in our estimates from the analysis of the different sampling cohorts. Ideally, our findings should be confirmed in longitudinal cohorts, with well-recorded past exposure and vaccination histories and subsequent paired assessment of immunological measurements and susceptibility to influenza infection and disease.

Our study provides an immunological link between birth year and influenza susceptibility. Understanding the immunological basis of birth-cohort effects is critical in appreciating the epidemiological and evolutionary impact of such phenomena, as well as in identifying obstacles that should be overcome and strategies that leverage the protective effects of imprinting. Our finding of birth-cohort specific biases in HI reactivity opens up the possibility to understand how immune pressure from differentially imprinted cohorts may shape the evolutionary trajectory of antigenically evolving viruses. Further immunological readouts that may identify individuals at higher risk of infection with a specific influenza subtype or lineage may also assist in the design of optimal cohort-specific intervention strategies to protect against future outbreaks.

## Acknowledgements

We thank the participants for their generous involvement and provision of samples. We are grateful for the CSL investigators that originally initiated this study NCT00959049 trial. We are grateful to Marcos Vieira, Katelyn Gostic and Sarah Cobey for discussions and advice on data analysis. The work has been generously supported by the Morningside Foundation and by Australian National Health and Medical Research Council Investigator grants (M.K., A.K.W., J.A.J and S.J.K.). The WHO Collaborating Center for Reference and Research on Influenza is supported by the Australian Government Department of Health.

## Author contributions

M.K. designed and supervised the study. M.K., L.S.U.S., M.A., N.S. performed the experiments. M.K., P.E., L.S.U.S., M.W., N.S., Y-M.D. and D.P. analysed data; M.A.C, T.M.R, J.A.J, S.R., S.J.K, A.K.W and I.G.B. provided samples, reagents and/or data critical for the study. M.K., J.A.J, A.KW, P.E. & D.P. contributed to drafting of the manuscript. All authors reviewed the final version of the manuscript.

## Methods

### Cohorts and serology datasets

The main dataset of this study was generated using cross-sectional serum samples collected and stored between 1992 and 2020. Samples were collected prior to any recorded influenza vaccination in that year. Samples from adults were collected under study protocols that were approved by the University of Melbourne Human Research Ethics Committee (2056689) or were provided by the WHO Collaborating Centre for Reference and Research on Influenza in Melbourne (WHOCCRRI). Human clinical samples that are supplied to the WHOCCRRI fall under the terms of reference of the WHO Global Influenza Surveillance and Response System (GISRS (43)) and the use of these samples for influenza vaccine development is permitted without additional ethics approval being held by WHOCCRRI. Samples from children aged 1-18 years old in 2009 were previously collected as part of a clinical trial NCT00959049 (44). All participants provided written informed consent in accordance with the Declaration of Helsinki. In addition, we analysed a previously published data set(23) of 84 adults sampled in 2014 and titrated HI assays against IBV isolates from the three lineages. We additionally analysed existing HI data from 1044 cross-sectional samples from 2000-2020 which had been titrated against the vaccine strains of the year the samples were collected at the WHOCCRRI.

Birth cohort groups were defined based on combined assessment of (i) the probability of most-likely lineage of first infection described by Vieria *et al*. (7); (ii) the previously reported phylogenetics and antigenic properties of IBV isolates described (11, 12) and (iii) the previously reported IBV seroprevalence by age group (45, 46, 47). Specifically, individuals born between 1981-1998 had at least a 50% probability of B/Yamagata imprinting (7), exhibit higher HI to B/Yamagata isolates than other lineages (Figure 1B and S3) and virus isolates from that time period belong to the B/Yamagata lineage (11, 12). Individuals born from 1999 onwards had equal probabilities of imprinting by either lineage or being naïve (7) and were grouped into the 1999-2008 (latest birth year of dataset) cohort. Individuals born between in 1980 or earlier had at least a 50% probability of being imprinted by a non-Yamagata lineage. As the earliest isolate described to have antigenic similarity to the B/Victoria lineage is from 1972 (12) (Figure 1B and S3) and IBV seroprevalence increases for the first 10-15 years of life (45, 46, 47), we chose 1961-1980 as the cohort with B/Victoria imprinting and individuals born prior to 1960 as those with Ancestral IBV imprinting. As there is no antigenic or sequence information about IBV isolates prior to 1940, and HI data from Figure 1B and S3 indicated that viruses prior to 1940 maybe differ from those circulating after 1940, we split individuals born prior to 1961 into the 1917 (earliest birth year of the dataset) – 1939 birth cohort, and the 1940 (first IBV isolate)-1960 birth cohort. We iterate that the birth year boundaries chosen here should not be considered as absolutes. The discrete birth cohorts are used to facilitate analyses of additional variables to complement analyses of the effects of birth year as a continuous variable.

### Viruses and reagents

Details of influenza B isolates used in this study are provided in Figure S1A. Egg isolates were propagated in 10-12 day old embryonated chicken eggs. Cell isolates were propagated in Madin Darby Canine Kidney (MDCK) cells in the presence of TPCK-treated trypsin. The human monoclonal antibodies used for antigenic characterisation have been previously described (CR8033(21), C12G6(22) and mAb29, mAb47, mAb15 (20)). Viruses were sequenced via Sanger sequencing and aligned using the multiple alignment using fast Fourier transform algorithm in MegAlign Pro 13 (DNASTAR Lasergene 13). A time-structured phylogenetic tree was estimated using BEAST 1.10.4 (48). Prior to the BEAST analysis an ML tree was used to determine the degree of clock-like behaviour of each data set by performing linear regressions of root-to-tip distances against year of sampling, using TempEst (49). Time-stamped data were analysed under the uncorrelated lognormal relaxed molecular clock (50), and the SRD06 codon-structured nucleotide substitution model (51). The Bayesian skyline coalescent tree prior was used. We compared log marginal likelihood following steppingstone sampling of both strict and uncorrelated lognormal relaxed molecular clocks, as well as a constant size and Bayesian skyline coalescent and selected the best model. One hundred million generations were performed and convergence was assessed using Tracer v1.6 (http://tree.bio.ed.ac.uk/software/tracer/). Maximum credibility clade trees were generated using TreeAnnotator v1.8. Phylogenetic trees were visualized in FigTree v1.4.4 (http://tree.bio.ed.ac.uk/software/figtree/).

### HI assays

Serological analysis by HI was performed according to the WHO Global Influenza Surveillance Network protocols (52), with the exception that volumes were reduced to 25 μL of sera, virus (4 HA units) and 1% turkey erythrocytes (0.33% final concertation). Human sera were treated with receptor destroying enzyme (Denka Sieken) and adsorbed with 5% erythrocytes prior to testing. For each individual, all titrations were performed using the same batch of erythrocytes. Samples were tested over two-fold serial dilutions from 1:10 to 1:20,480. IBV isolate were ether treated according to WHO Global Influenza Surveillance Network protocols (52). Monoclonal antibodies with known reactivity patterns were run with each new batch of samples/erythrocytes and were accepted if HI titres were within two-fold of expected values.

### Microneutralization assay

The neutralisation activity of serum was examined using a microneutralisation assay. Briefly, MDCK cells were seeded in 96-well plates at 3 x 10^4^ per well one day prior to the assay and incubated at 37°C overnight. Serum samples were heat inactivated at 56°C for 30 min and serially diluted (2-fold, starting at 1:10) in Virus Infection Media (VIM - DMEM supplemented with PSG, 1 mM Sodium pyruvate, 0.5 % BSA and HEPES). Serum samples were incubated for 60 min at 37°C with an equivalent volume of 100 TCID virus diluted in VIM. After incubation, MDCK cells were washed twice in PBS. VIM (100μL) supplemented with 3 μg/mL TPCK-treated trypsin and 100 μL virus-serum mix were added to the MDCK monolayer. Control wells of virus alone and VIM alone were included on each plate. Virus input was back-titrated on MDCK cells. Cells were incubated for 3 days at 37°C and the presence of virus was determined by haemagglutination assay. Briefly, 25 μL supernatant from each well were mixed with 25 μL of 1% (v/v) Turkey erythrocytes, incubated for 30 min at room temperature and presence of virus was recorded. Microneutralisation titres were determined using the reciprocal of the highest dilution at which virus neutralisation was observed.

### Statistical analysis

Distributions of medically attended IBV cases by birth year were determined by accessing data from the Global Initiative on Sharing All Influenza Data (GISAID) database (29^th^ August 2022) and grouping based on reported lineage. Distributions were determined from cases reporting the age of the host from which birth year was calculated. Descriptive statistics of HI for different isolates and birth cohort are provided in Tables S1-6 as they relate to different figures. P-values were generated from a Kruskal-Wallis test with Dunn’s correction for multiple comparisons, Friedman’s test with Dunn’s correction for multiple comparisons or a Wilcoxon matched-pairs signed rank test as appropriate, using GraphPad Prism 9.5.0. To construct and compare antibody landscapes across strains, we used GAMs fit to log_2_ titres against birth year or age at virus isolation. Plots were generated with ggplot2 (53). We used the GAM function from the R package mgcv and accounted for repeated measurements on each individual through specification of a random effect.

### Estimating the probability of infection from HI data

We use the following equation as per Coudeville et al.(24) to represent the probability of infection (equation 1):

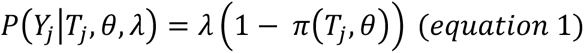

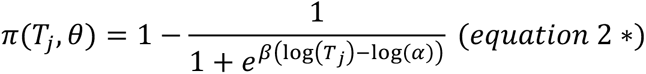

Where alpha and beta are parameters of the function which defines the contribution of HI titres to protection (referred to as the HI protection curve in Coudeville et al.) (equation 2). Alpha is closely linked to the 50% protection titre and beta is related to the slope of the protection curve. Lambda is the baseline risk of infection in the absence of any HI-related protection. Coudeville et al. assume the alpha and beta parameters are log normally and normally distributed, respectively, 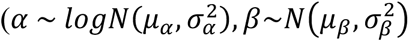, and provide estimates for the mean and standard deviation for each of the parameters with 95% confidence intervals. From these we specified a reasonable, approximate, distribution for each parameter that we were able to sample from (Table 1)

**Table 1:** The estimates used from the Coudeville paper for the equation parameters and approximate distribution we used to sample from.

| Parameter | Coudeville estimate [95% CI] | Approximate distribution |
| --- | --- | --- |
| Mean $\mu_\alpha$ | 2.844 [ 2.25, 3.36] | N(2.844, 0.08) |
| Standard deviation $\sigma_\alpha$ | 0.845 [0.44, 1.44] | logN(-0.213, 0.088) |
| Mean $\mu_\beta$ | 1.299 [1, 1.69] | logN(0.253, 0.018) |
| Standard deviation $\sigma_\beta$ | 0.376 [0.1, 0.76] | logN(-1.112, 0.268) |
| Lambda $E[\lambda_i]$ | 0.482 [0.41; 0.57] | logN(-0.733, 0.007) |

We sampled from these distributions for alpha and beta to re-create the HI protection curve shown in Coudeville et al. (equation 2 – note the HI protection curve function was slightly modified to ensure samples taken from the 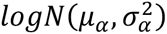 are on the equivalent scale to log(HI) values). The HI protection curve corresponding to our sampled parameters was not an exact fit to the protection curve reported in Coudeville et al., which we believe may be due to their results allowing for variability in the observed HI titres. However, it does fit within the 95% confidence limits and for the purposes of this study, the protection curve has been used to compare the probability of infection curves, and therefore should not be used as absolute probability of infection.

To establish the ‘probability of infection’ curves we sampled 10,000 values from each alpha, beta and lambda parameter distribution and applied these to 10,000 sampled HI titres from the GAM models. These values were used in the above equation and median and IQR values for the probability of infection were calculated.

## Data availability

The raw HI data will be provided as a supplementary table following acceptance of the manuscript. All other data and code used for analyses are available from the corresponding author upon reasonable request.

---

**Figure S1.**
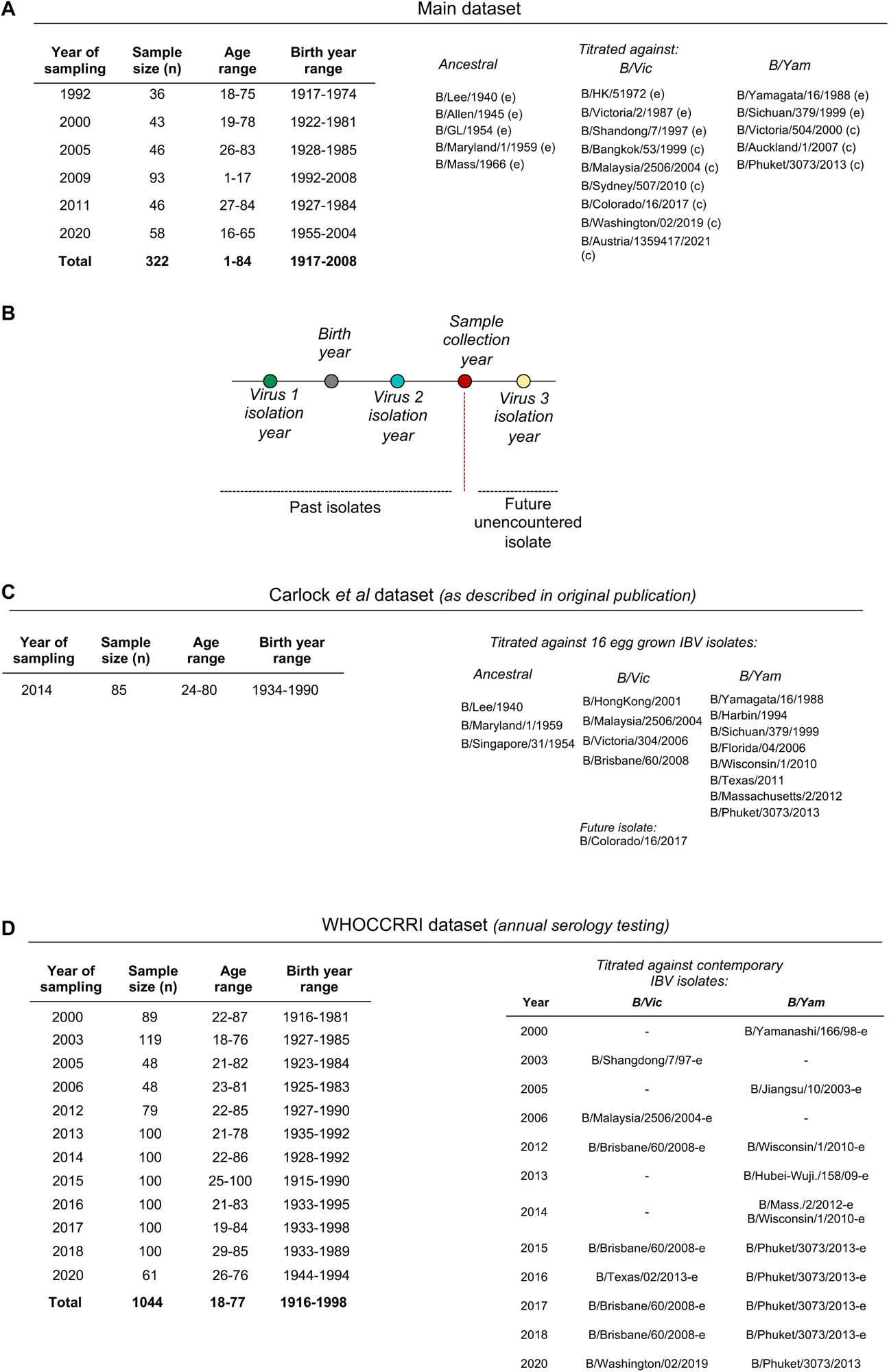
Cohort and dataset details. **(A)** Details of the cohort associated with the main dataset generated for this study. The virus passage is indicated for each isolate (e – egg; c – cell). **(B)** Schematic of the relationships between year of virus isolation, birth year and sampling year. **(C)** Details of the cohort associated with the dataset generated by Carlock et al. **(D)** Details of the cohort associated with the WHOCCRI dataset.

**Figure S2.**
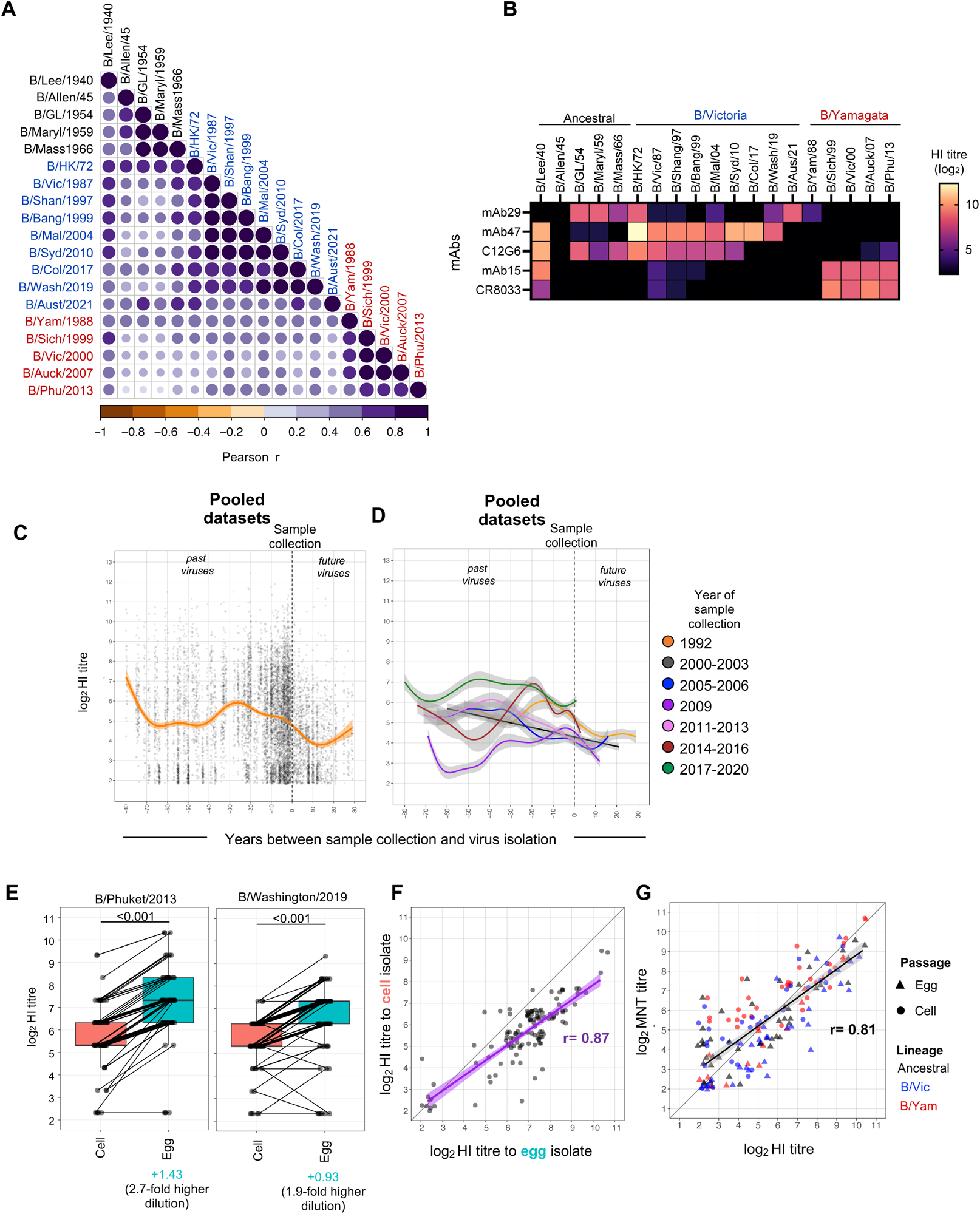
Overview of HI titres. **(A)** Correlations between antibody titres against the 19 IBV isolates from the main dataset (n=322 serum samples). Virus isolates have been ordered by hierarchical clustering. **(B)** Antigenic characterization using human monoclonal antibodies against the IBV HA. End point titres from an HI assay are shown starting at 50μg/ml. **(C)** Log_2_ HI titres against time (years) between sample collection and virus isolation (across all three datasets). **(D)** Log_2_ HI titres against time (years) between sample collection and virus isolation (across all three datasets) with individuals pooled into sample collection groups. **(E)** Comparison of HI titres against cell and egg isolates in paired serums samples tested against cell or egg grown B/Phuket/2013 and B/Washington/2019. The number in blue indicated the difference in titres on the log_2_ scale. P-values were generated from a Wilcoxon matched-pairs signed rank test (n=61 samples from 2020). **(F)** Correlation between log_2_ HI titres against cell and egg grown B/Phuket/2013 and B/Washington/2019 viruses as described in C. **(G)** Correlation between log_2_ HI titres and microneutralization titres in 17 individuals tested against 10 viruses.

**Figure S3.**
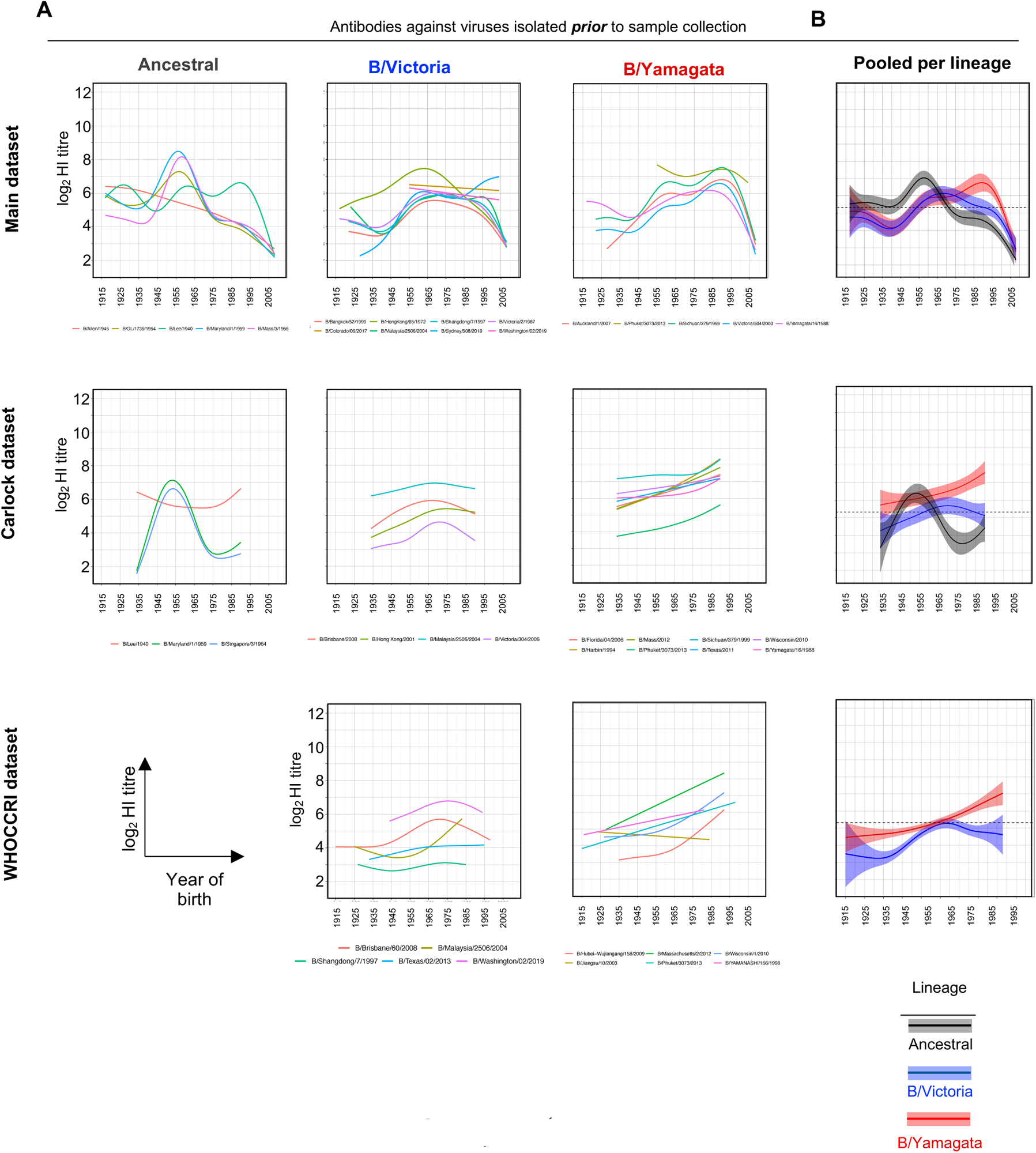
HI titres to past isolates against year of birth. **(A)** Log_2_ HI titres to different isolates against year of birth separated by IBV lineage and dataset. The lines represent estimated mean HI titres from generalized additive models (GAMs). **(B)** Estimated mean HI titres against past isolates from each lineage that circulated prior to sample collection, against the year of birth for each participant. The lines represent the estimated mean HI titre from generalized additive models (GAMs) with shading representing 95% CIs, accounting for repeated measurements on each individual by specifying a random effect.

**Figure S4.**
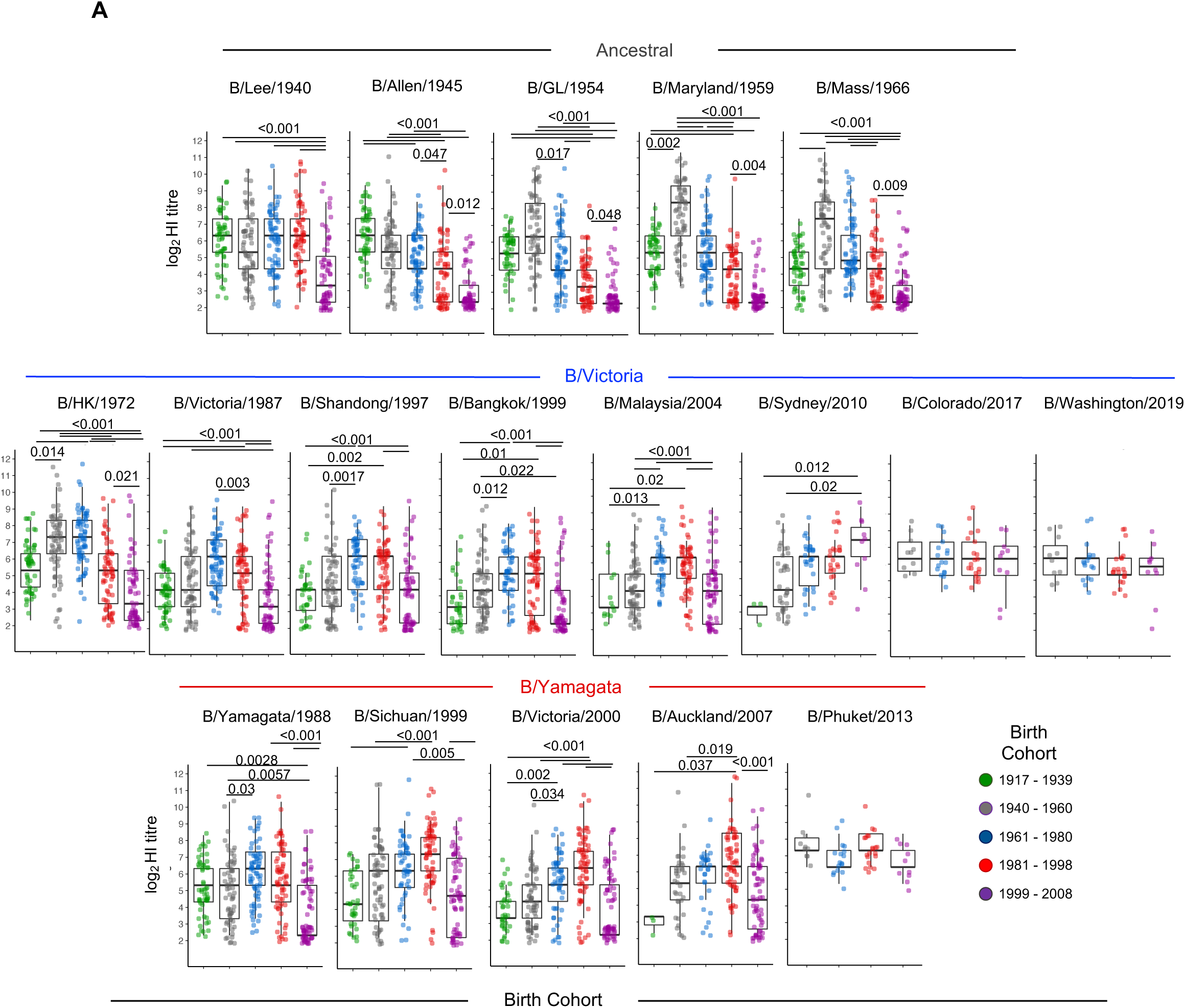
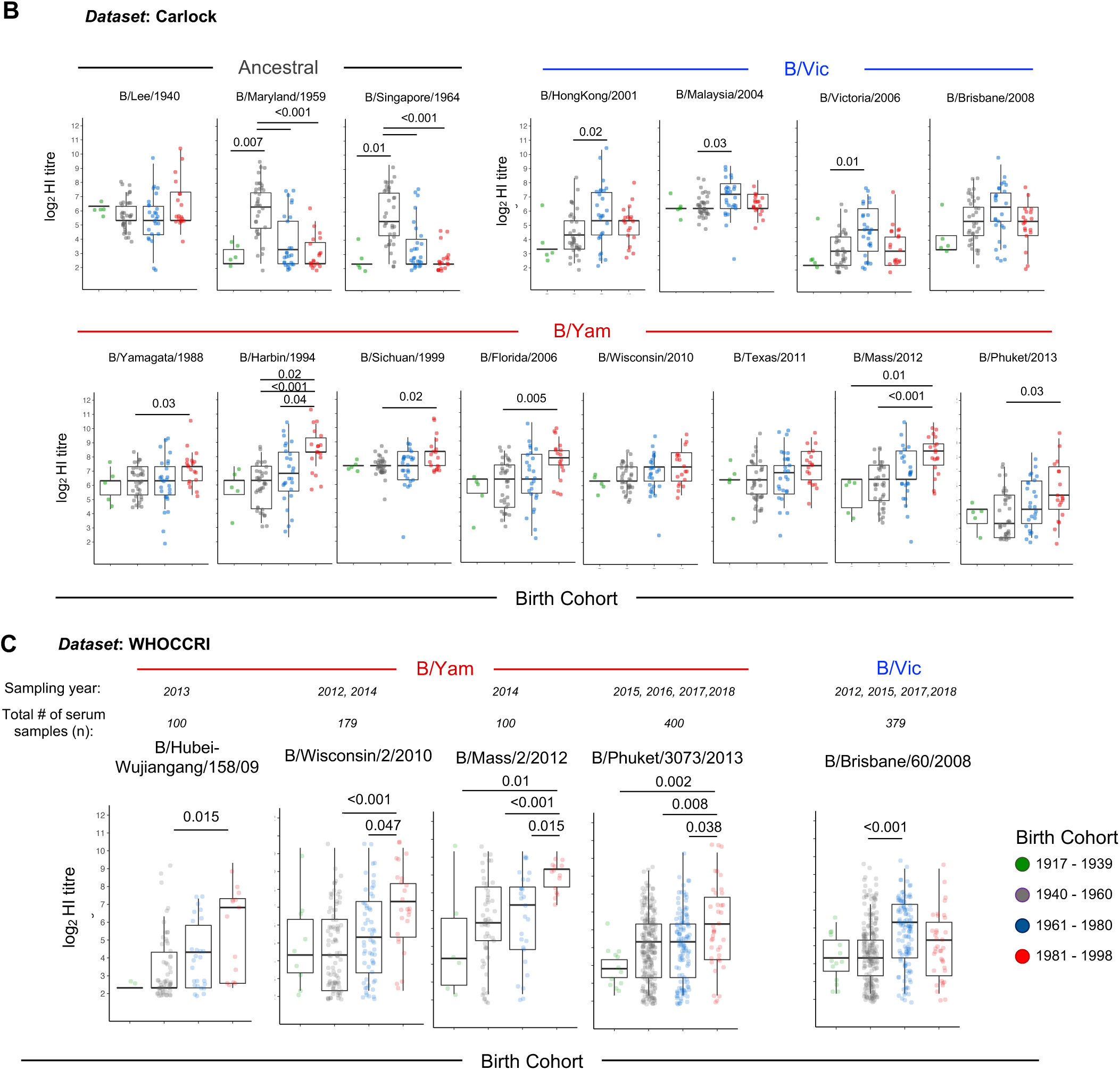
HI reactivity against past isolates by birth cohort across different datasets. Box-plots of antibody titres to specific IBV isolates for each birth cohort for **(A)** the main dataset, **(B)** the Carlock et al dataset and **(C)** the WHOCCRI dataset. P-values were generated frmo a Kruskal-Wallis test with Dunn’s correction for multiple comparisons.

**Figure S5.**
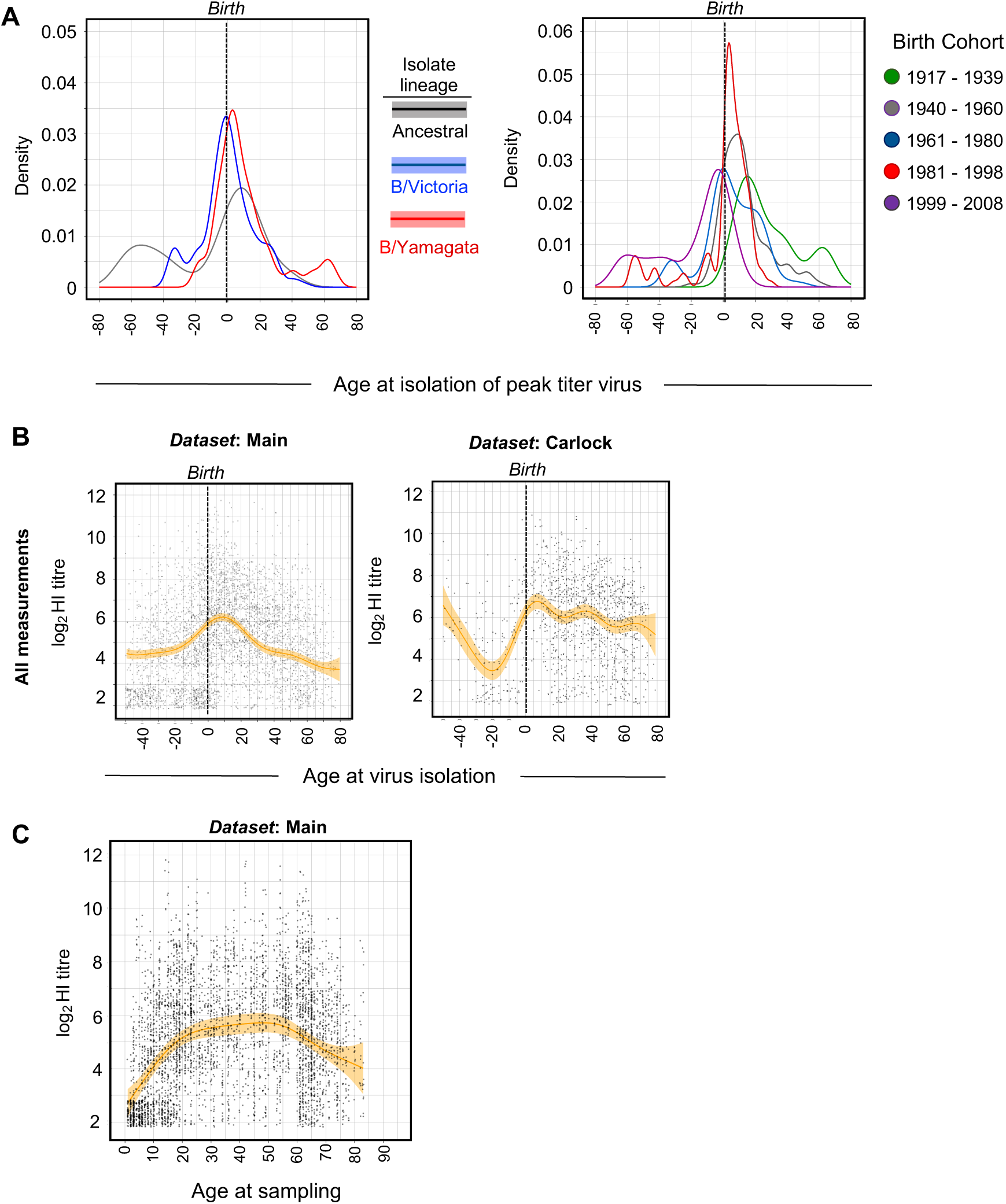
Antigenic seniority in IBV HI titres. **(A)** Distribution of the age of the participant at the time of isolation of the strain to which they had highest antibody titres, grouped by IBV lineage or birth cohort. **(B)** HI titres relative to the age of the participant at the time of virus isolation. **(C)** HI titres relative to the age of the participant at the time of sampling. For (B) and (C) the lines represent estimated mean HI titres from generalized additive models (GAMs) with shaded region representing 95% CIs, accounting for repeated measurements on each individual by specifying a random effect. Only titrations of viruses isolated prior to sample collection are included. Analysis shown for measurements from the main (A, B, C) or Carclock et al (B) datasets.

**Figure S6.**
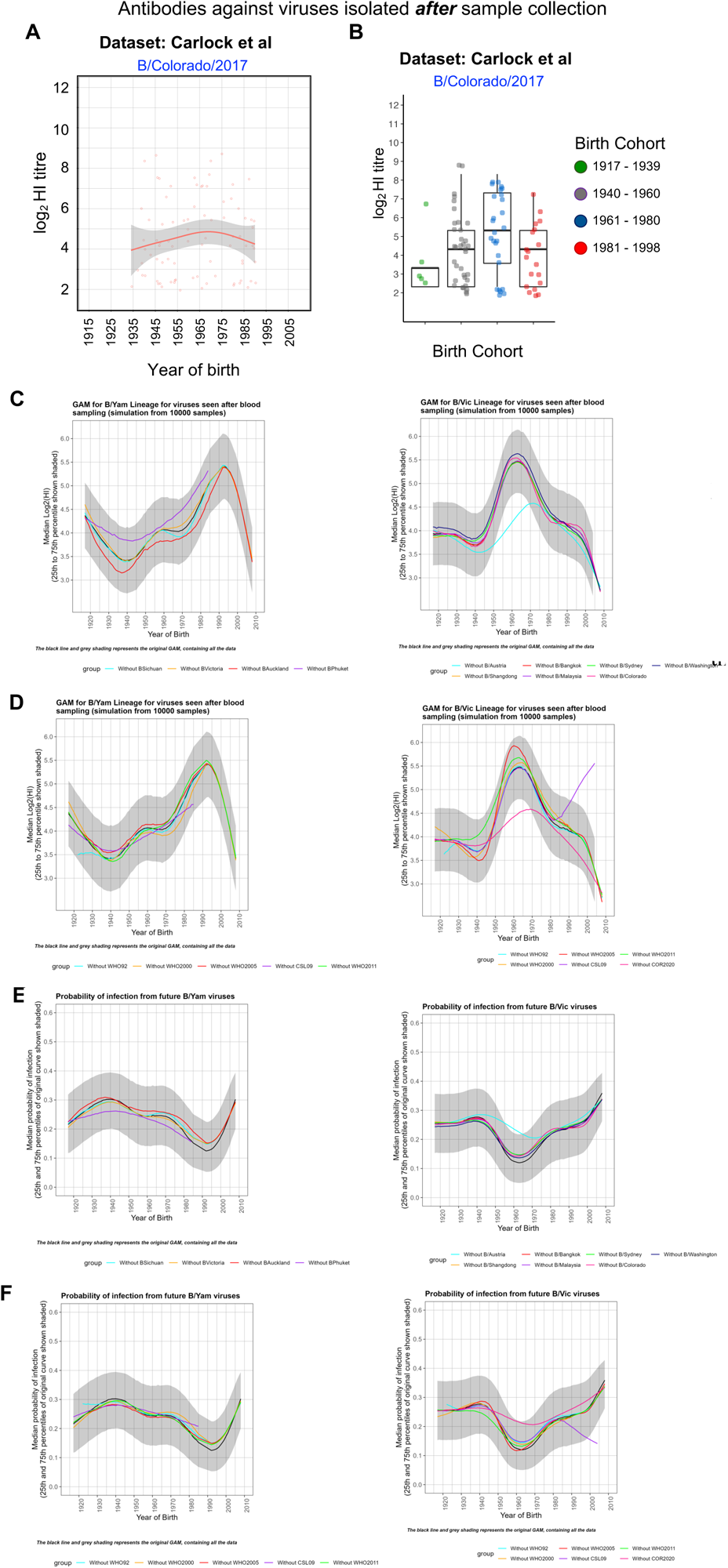
HI titres and estimated susceptibility to future isolates varies by birth year. **(A)** Estimated mean HI titres against the future unencountered B/Colorado/02/2017 from the Carlock et al dataset. **(B)** Box-plots of HI titres to future unencountered B/Colorado/02/2017 for each birth cohort from the Carlock et al dataset. **(C-F)** Sensitivity analysis of estimated HI (C-D) and probabilities of infection (E-F) by birth year for the effects of different virus isolates (C, E) or sampling year groups (D, F). The median estimated HI or probability is shown and the shaded areas represent the 25^th^ and 75^th^ percentiles.

